# Close agreement between deterministic vs. stochastic modeling of first-passage time to vesicle fusion

**DOI:** 10.1101/2021.07.30.454536

**Authors:** Victor Matveev

## Abstract

Ca^2+^-dependent cell processes such as neurotransmitter or endocrine vesicle fusion are inherently stochastic due to large fluctuations in Ca^2+^ channel gating, Ca^2+^ diffusion and Ca^2+^ binding to buffers and target sensors. However, prior studies revealed closer-than-expected agreement between deterministic and stochastic simulations of Ca^2+^ diffusion, buffering and sensing, as long as Ca^2+^ channel gating is not Ca^2+^-dependent. To understand this result more fully, we present a comparative study complementing prior work, focusing on Ca^2+^ dynamics downstream of Ca^2+^ channel gating. Specifically, we compare deterministic (mean-field / mass-action) and stochastic simulations of vesicle exocytosis latency, quantified by the probability density of the first-passage time (FPT) to the Ca^2+^-bound state of a vesicle fusion sensor, following a brief Ca^2+^ current pulse. We show that under physiological constraints, the discrepancy between FPT densities obtained using the two approaches remains small even if as few as ∼50 Ca^2+^ ions enter per single channel-vesicle release unit. Using a reduced two-compartment model for ease of analysis, we illustrate how this close agreement arises from the smallness of *correlations* between fluctuations of the reactant molecule numbers, despite the large magnitude of the fluctuation *amplitudes*. This holds if all relevant reactions are *heteroreaction* between molecules of different species, as is the case for the bimolecular Ca^2+^ binding to buffers and downstream sensor targets. In this case diffusion and buffering effectively decorrelate the state of the Ca^2+^ sensor from local Ca^2+^ fluctuations. Thus, fluctuations in the Ca^2+^ sensor’s state underlying the FPT distribution are only weakly affected by the fluctuations in the local Ca^2+^ concentration around its average, deterministically computable value.

**Statement of Significance:** Many fundamental Ca^2+^-dependent cell processes are triggered by local Ca^2+^ elevations involving only a few hundred Ca^2+^ ions. Therefore, one expects large Ca^2+^ concentration fluctuations, which are ignored by deterministic reaction-diffusion modeling approaches. However, more accurate stochastic approaches require tracking trajectories of individual Ca^2+^ ions and its binding targets, which is very computationally expensive. This study reveals conditions under which Ca^2+^-dependent processes like secretory vesicle fusion can be modeled using efficient deterministic approaches, without sacrificing significant accuracy. We find that deterministic methods can accurately predict the delay to the fusion of a neurotransmitter-containing vesicle, as long as the number of Ca^2+^ ions is above about 50. We reveal factors that explain the limited impact of stochastic fluctuations in this case.

## I. INTRODUCTION

Many fundamental cell processes such as myocyte contraction and synaptic and endocrine secretory vesicle fusion are controlled by highly localized Ca^2+^ signals resulting from the opening of transmembrane Ca^2+^ channels [1-5]. Imaging local Ca^2+^ concentration required for the study of these processes is a great challenge due to physical limitations on the spatial and temporal resolution of optical imaging, and necessarily perturb the Ca^2+^ signals being measured. Therefore, Ca^2+^ modeling continues to play a crucial role in the study of a variety of Ca^2+^-dependent cell mechanisms. The main decision facing a modeler is whether to choose a deterministic or a stochastic solver [5-10]. While deterministic mass-action reaction-diffusion approach offers superior computational efficiency, it completely ignores local fluctuations in Ca^2+^ resulting from the stochastic Ca^2+^ channel gating, diffusion, and biochemical reactions. For processes controlled by single-channel Ca^2+^ domains, fluctuations are considerable: a typical Ca^2+^ current of 0.2 pA and duration of 0.2 ms translates to an influx of only about 120 Ca^2+^ ions, many of which become bound to mobile buffers and transmembrane proteins before reaching their downstream target sensors [11-13]. Thus, one should expect large Ca^2+^ concentration fluctuations at the location of a relevant Ca^2+^ sensor that only a few Ca^2+^ ions will reach [13, 14]. Stochastic fluctuations were shown to have functional consequences in reaction-diffusion models of a variety of cell processes (see e.g. [15-18]). For Ca^2+^-controlled processes, such stochastic effects are especially pronounced in the presence of Ca^2+^–induced Ca^2+^ release (CICR) responsible for excitation-contraction coupling in myocytes, since CICR introduces a direct feedback between Ca^2+^ fluctuations and Ca^2+^ influx [19-28].

Despite the widely recognized importance of stochastic effects, comparative studies suggest that downstream of stochastic channel gating, the discrepancy between deterministic and stochastic simulations of Ca^2+^ diffusion, buffering and binding can be surprisingly small [24, 29]. Therefore, in the absence of CICR, computationally inexpensive simulations of Markovian stochastic channel gating can be combined with deterministic models of Ca^2+^ diffusion and binding (either compartment-based or spatially resolved), leading to computationally inexpensive methods that avoid simulations of particle-based Brownian motion and stochastic reactions [7, 27-30]. This in fact has often been the approach even in the modeling of CICR, where Ca^2+^ channel gating is the primary source of fluctuations [5, 19-21, 31-39]. This simplified approach has also proved useful in the study of vesicle fusion [40, 41], Ca^2+^ signaling in dendrites [42], and Ca^2+^- dependent K^+^ channels [43].

Despite a large number of relevant studies, in particular a comprehensive comparative computational study by Modchang et al. [29], a deep understanding of the discrepancy between stochastic and deterministic simulations downstream of stochastic Ca^2+^ channel gating effects is still lacking. As a result, as has been noted previously [24], the choice between deterministic and stochastic solvers is usually made on a completely *ad hoc* basis. In particular, it is widely accepted that deterministic reaction-diffusion methods are highly inaccurate in the modeling of any biochemical process involving a small number of molecules (e.g. the number of Ca^2+^ ions, *N*_Ca_). This expected inaccuracy is due to the well-known discrepancy between mass-action and stochastic representations of any nonlinear process [44, 45].

However, here we show that the size of this discrepancy between deterministic and stochastic approaches is not always as large as expected from an often-used naïve 1/√*N*_ca_ scaling, and depends not only on the size of the Ca^2+^ fluctuations, but also on the type of observable targeted by the modeling, and the type of reactions involved. In particular, following several recent studies, we focus on the first passage time distribution (FPTD) to full finding of a model Ca^2+^ sensor for vesicle fusion in the presence of bimolecular Ca^2+^ buffering reactions [13, 14, 46-51]. We consider a maximally reduced but spatially resolved model that contain only two sources of fluctuations: (1) the diffusive fluctuations in Ca^2+^ concentration, and (2) the fluctuations due to Ca^2+^ binding and unbinding to Ca^2+^ buffers and sensors. Considering such a reduced model reveals more clearly the interplay between fluctuations due to diffusion and reaction in limiting the accuracy of the mean-field modeling of Ca^2+^ sensor binding time. Our main finding is that the discrepancy between deterministic and stochastic estimates of FPTD can be negligible as long as the number of Ca^2+^ ions is above the threshold of about ∼50 (see Figs. 1-3). Finally, to clarify our conclusions, we will also analyze an even simpler two-compartment model of Ca^2+^ diffusion and binding, following the approach of G.D. Smith and S.H. Weinberg [26, 48, 52]. Our results further elucidates how the interplay between the diffusion and the reaction time scales affects the accuracy of the deterministic approach [15, 23, 24, 29, 38, 48].

**Figure 1.**
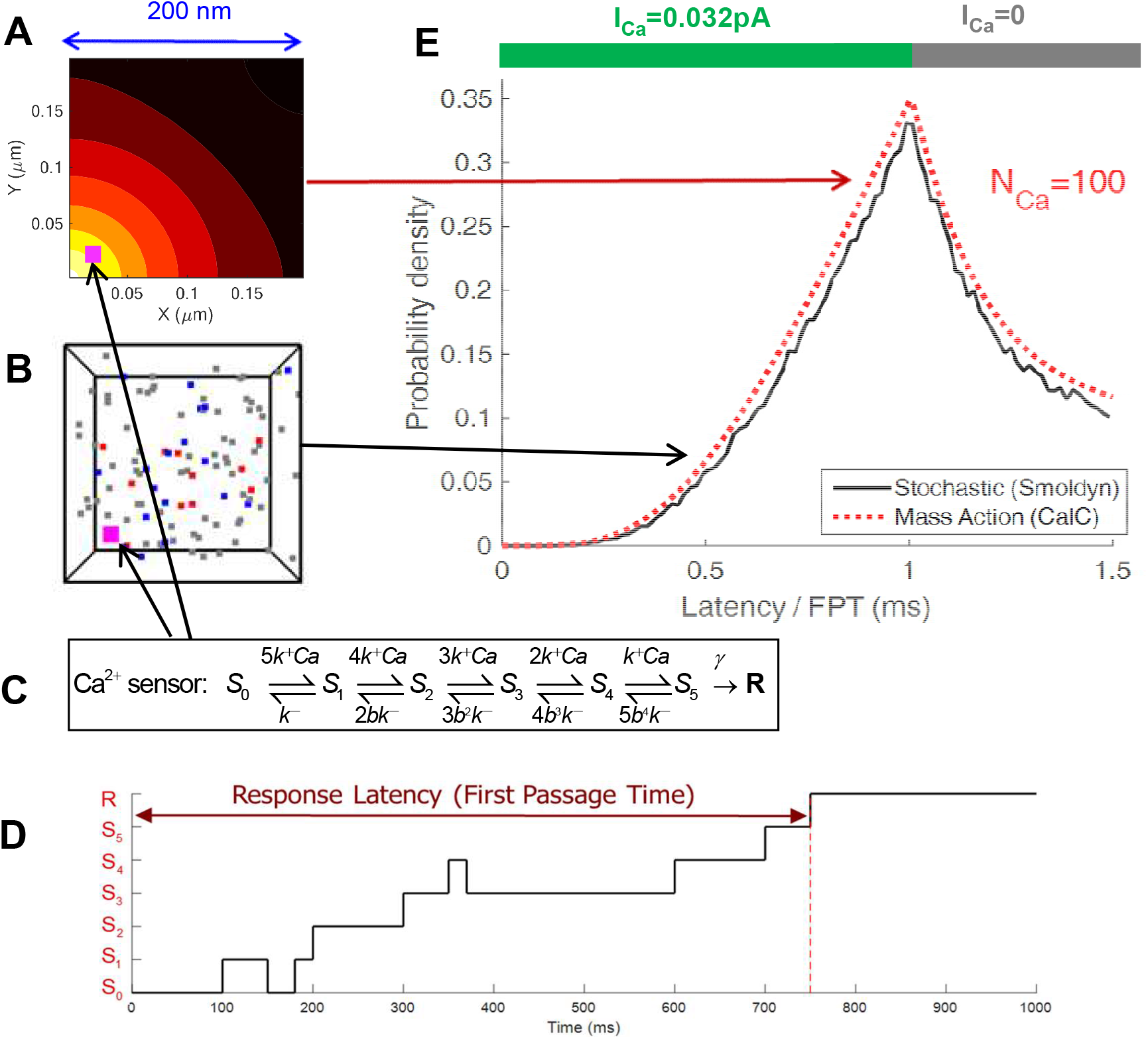
Deterministic vs. stochastic simulation of buffered Ca^2+^ diffusion and binding to a stationary sensor. Ca^2+^ and buffer molecules diffuse within a (0.2μm)^3^ cube, with diffusivities *D*_Ca_=0.2μm^2^/ms and *D*_B_=0.05μm^2^/ms, respectively. Ca^2+^ sensor is 33nm away from a point Ca^2+^ source (Ca^2+^ channel) in the corner of the cube. Ca^2+^ current pulse of 0.032 pA lasts 1ms, letting in a total of 100 Ca^2+^ ions. Buffer has a concentration of 20μM and affinity of 1μM, corresponding to a total of 96 ions. **(A)** One time frame of a 2D slice of the mass-action simulation of [Ca^2+^] on a color-coded logarithmic scale, 0.2ms after Ca^2+^ channel opening. **(B)** One time frame of the stochastic Smoldyn iteration shows the locations of all tracked particles 0.2ms after Ca^2+^ channel opening (*magenta square*: Ca^2+^ sensor; *red squares*: Ca^2+^ ions; *gray and blue squares*: free and bound buffer molecules, respectively). Symbol sizes are not to scale. (**C**) Stationary vesicle fusion sensor undergoes 5 Ca^2+^ binding steps with progressively decreasing unbinding rate, i.e. increasing binding cooperativity (Eq. 4) [56, 57], (**D**) Sensor state transition sequence illustrates a single MCMC trial of the stochastic Smoldyn simulation. FPT is computed from the start of the Ca^2+^ current pulse to the time of sensor transition to the final state ***R***. (**E**) Comparison of FPTD obtained from the histogram of stochastic trials shown in **D** (*black curve*), or from deterministic reaction-diffusion simulation (Eqs. 2-5) (*red dotted curve*), with binding depletion ignored (σ_S_=0 in Eq. 2).

**Figure 2.**
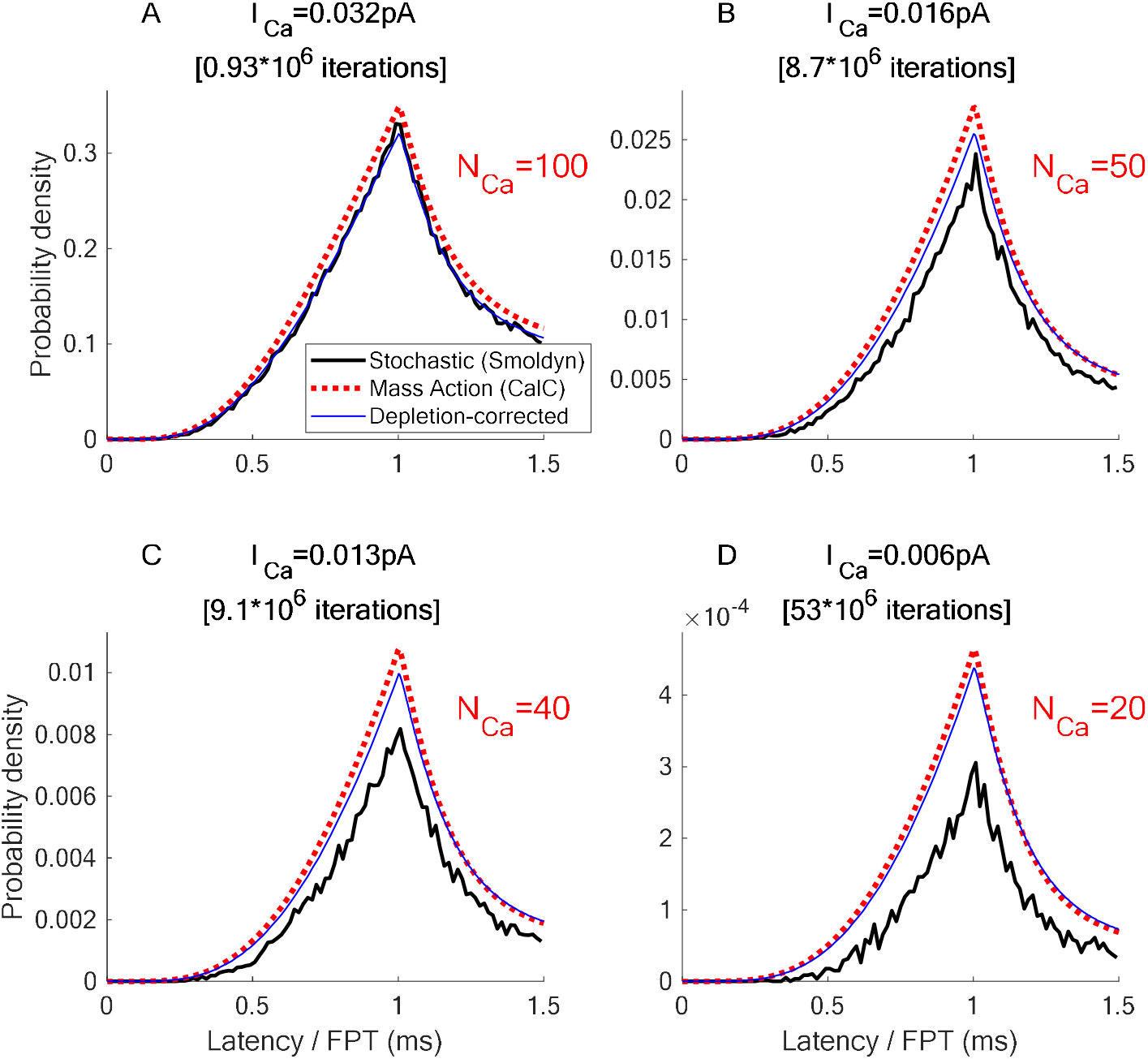
Mass-action vs stochastic estimation of FPTD in response to a 1ms-long Ca^2+^ current pulse of varying amplitudes. All parameters as in Fig. 1, except for the varying Ca^2+^ current values and the total number of Ca^2+^ ions entering during the 1-ms current pulse (*N*_Ca_), as indicated in text labels. In all panels, *black solid curves* represent the histograms of FPT to full sensor binding, the *dotted red* curves show the corresponding deterministic estimate of FPTD (Eq. 5), with binding depletion ignored (σ_S_=0 in Eq. 2), while the *solid blue* curves show the deterministic estimate of FPTD, with binding depletion taken into account (Eqs. 6-7). Note the difference in scale: the cumulative probability of full binding within 1.5 ms of channel opening is about 18% in *A* (N_Ca_=100), while in *D* (N_Ca_=20) it is on the order of 10^−4^, explaining the large number of MCMC trials required in the latter case, since most binding events happen in the long tail of this distribution.

**Figure 3.**
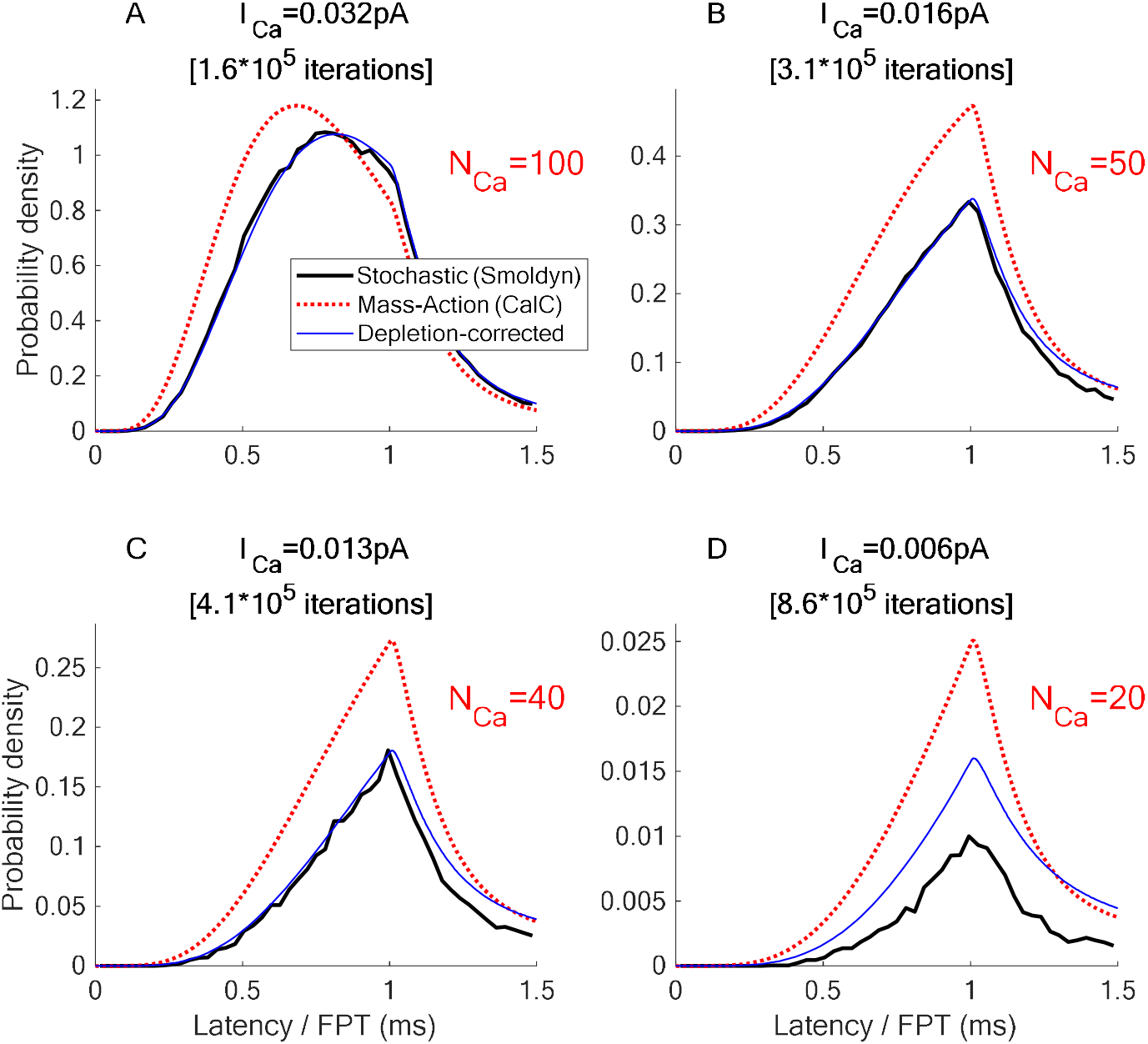
Mass-action vs stochastic simulation of FPTD for unphysiologically large sensor binding rate, and a greater distance between release sensor and the Ca^2+^ channel, in response to a 1ms-long Ca^2+^ current pulse of varying amplitude. Simulation parameters are similar to the ones in Figs. 1-2, except a factor of 10 faster exocytosis sensor binding rate *k*^+^, requiring a smaller MC simulation time step of Δ*t*=0.1μs, larger diffusion volume of (0.3μm)^3^, and larger distance from channel to sensor of 83 nm. The Ca^2+^ current value and the total number of Ca^2+^ ions entering during the 1-ms long current pulse is indicated in the panel titles. In all panels, *black solid curves* represent the histograms of FPT to full sensor binding, the *dotted red* curves show the corresponding deterministic estimates of FPTD (Eq. 5), with binding depletion ignored (σ_S_=0 in Eq. 2), while the *solid blue* curves show the deterministic estimate of FPTD, with binding depletion taken into account (Eqs. 6-7). Note the difference in scale: the cumulative probability of binding within 1.5ms is about 74% in *A* (N_Ca_=200), while in *D* (N_Ca_=20) it is ∼0.5%.

## II. METHODS

### II.1 Deterministic 3D mass-action / mean-field approach

Let us first describe the deterministic model of Ca^2+^-dependent exocytosis. For the sake of simplicity, we consider the case of a single dominant Ca^2+^ buffer with a single Ca^2+^ ion binding site, as described by the reaction

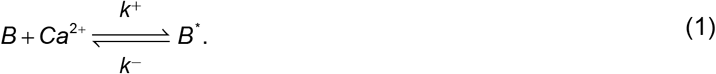

Here *B* and *B*^*^ represents the free buffer and Ca^2+^-bound buffer molecules, respectively (i.e. *B*^*\**^*=CaB*), and *k*^+^ (*k*^−^) are the Ca^2+^-buffer binding (unbinding) rates. Assuming isotropic diffusion and mass-action kinetics, this yields the following mass-action reaction-diffusion system [53-55]:

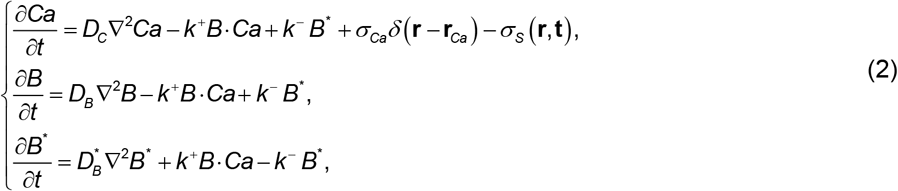

where *D*_*B*_, *D*^*\**^_*B*_ and *D*_C_ are the diffusivities of the free buffer, bound buffer and Ca^2+^, respectively, and *Ca, B* and *B*^*^ denote concentrations of Ca^2+^, free buffer and Ca^2+^-bound buffer, respectively. The delta function indicates a point source (channel) centered at location **r**_**Ca**_, with amplitude *σ*_Ca_ =*I*_**Ca**_/(2*F*) where *I*_**Ca**_ is the Ca^2+^ current strength, and *F* is the Faraday constant. The sink term *σ*_S_ describes Ca^2+^ binding to the model exocytosis sensor, as will be explained further below (see Eqs. 6-7).

We take a simple cube as the diffusion domain, with zero flux boundary conditions for Ca^2+^ and buffer on all boundaries. Reflective boundary conditions are well suited for simulating an array of Ca^2+^ channels on a large section of a flat cell membrane. Further, reflective boundary conditions ensure non-negligible probability of Ca^2+^ binding to a model exocytosis Ca^2+^ sensor (see below) within a short time after the channel opening, even if only 20 Ca^2+^ ions enter the model volume during the 1ms-long current pulse (see Figs. 1-3). Replacing no-flux conditions with Robin boundary conditions simulating Ca^2+^ pumps and exchangers on the part of the boundary containing the channel would be more realistic, but would introduce the same type of bimolecular Ca^2+^-binding reactions already present in this model, and therefore would not substantially impact the comparison of stochastic and deterministic simulation results.

The reaction-diffusion system given by Eqs. 2 is locally coupled to reactions describing a single stationary Ca^2+^ binding exocytosis sensor, with parameters inferred from the studies of neurotransmitter release at the calyx of Held synapse [56, 57]:

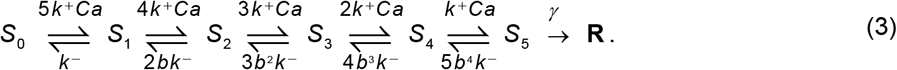

The rate parameters are *k*^−^=8.43 ms^−1^; *k*^+^=0.116 μM^−1^ms^−1^; *b*=0.25; γ=7.0 ms^−1^ (except Fig. 3, where the binding rate *k*^+^ is increased by a factor of 10). This reaction is converted to a system of deterministic ODEs describing the Markovian transitions between distinct sensor states:

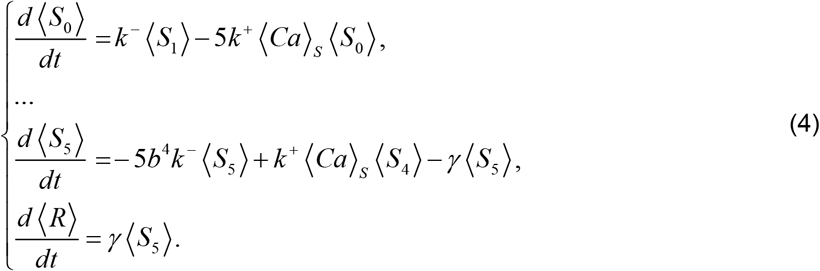

Here ⟨Ca⟩_S_ is the time-dependent [Ca^2+^] averaged over a small spherical volume describing the sensor, obtained by numerically solving Eq. 2, whereas ⟨*S*_1..5_⟩ and ⟨*R*⟩ are the time-dependent occupancy probabilities of the distinct states of the single-copy sensor. In this simplified approach, all rates in Eq. 4 are deterministic, with the forward binding rates proportional to the time-dependent but deterministic local Ca^2+^ concentration, ⟨Ca⟩_S_. Thus, Eq. 4 represents a mean-field description, neglecting stochasticity in the binding rates caused by local Ca^2+^ fluctuations. Nevertheless, this deterministic mean-field approach allows to estimate the fluctuation in the FPT to the fully-bound state *R* by computing its probability density, given by the transition rate to this final absorbing state [58]:

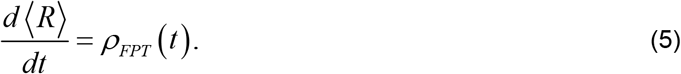

The primary goal of this study is to compare the FPT density (FPTD) given by Eq. 5 with the one obtained using fully stochastic simulation explained in the next subsection (see Figs. 1-3), in order to reveal the quantitative impact of various fluctuation sources on the latency to vesicle release.

In the simplest deterministic implementations of Ca^2+^ signaling models, the sink term in Eq. 2 is usually ignored, and therefore Ca^2+^ is not fully conserved. However, the binding of Ca^2+^ to exocytosis sensor can become significant when the Ca^2+^ influx current is small. As has been pointed out previously [24, 29], failure to account for the binding-induced Ca^2+^ depletion is not an inherent deficiency of the deterministic approach, and can be corrected by restoring full Ca^2+^ conservation. In our implementation, this is achieved by adding a localized Ca^2+^ sink term in Eqs. 2:

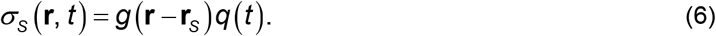

Here *g*(**r**−**r**_S_) has a finite support defined by the sensor volume, centered at the sensor location **r**_S_, with a total volume integral of 1, whereas *q*(*t*) is the total Ca^2+^ flux induced by the sensor reactions in Eq. 4:

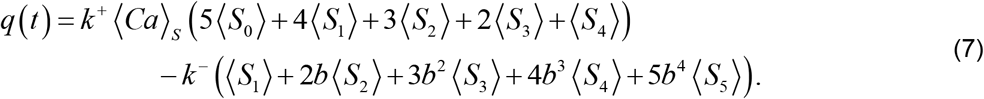

In our model, the sensor position **r**_S_ in Eq. 6 is close to the channel location **r**_Ca_, as shown in Fig. 1A,B. The distance between the two is either 33 nm (Figs. 1-2) or 83 nm (Fig. 3). The full deterministic simulation system is described by Eqs. 2-7, and satisfies exact Ca^2+^ conservation. However, in order to fully quantify the impact of this binding-induced Ca^2+^ depletion, we will repeat all simulations with and without the σ_S_ sink term in Eq. 2.

Note that Eqs. 2-7 describe the Ca^2+^ sensor as a *volume reactivity* (Doi) model, whereby the binding reaction takes place within a certain finite volume at a given time-dependent rate. In contrast, we will treat the same sensor binding as a *surface reactivity* (Smoluchowski) model in the stochastic simulation described in the next subsection, whereby the reaction takes place with certainty when the Ca^2+^ ion is within the binding distance of the sensor [59, 60]. However, the two sets of results presented below are nevertheless remarkably close to each other (see Figs. 1-3), with the sensor’s binding radius (the support of *g*(**r**) in Eq. 6) constituting one free matching parameter, set slightly larger than the Smoluchowski binding radius in the stochastic approach. We note that the exact Ca^2+^ conservation is exactly satisfied for any value of this parameter. In principle this free parameter could be avoided by using the surface reactivity model in the deterministic approach as well, but at the expense of greater complexity of implementing the prescribed flux given by Eq. 7 as a time-dependent boundary condition on the sensor’s surface.

Biologically, exocytosis sensors are positioned around the base of a vesicle, the latter acting as an obstacle to diffusion [13]. However, we deliberately excluded vesicles and other obstacles from our comparative analysis, since they would not contribute to the discrepancy between deterministic and stochastic simulations arising from the neglect of local Ca^2+^ fluctuations, unless molecular crowding effects are taken into account. The effect of diffusion obstacles on exocytosis has already been explored in detail [13, 46, 61-66].

All deterministic reaction-diffusion simulations are performed using the CalC (“Calcium Calculator”) software [67, 68], and take less than a minute of single-thread CPU time.

### II.2 Stochastic approach

Stochastic approach allows a more realistic modeling of the biochemical pathways leading to vesicle release, taking into account relevant fluctuations, albeit at the computational expense of repeated trials of an appropriate Markov Chain Monte Carlo (MCMC) simulation. However, accurately combining diffusion with second-order bi-molecular reactions described by Eqs. 3 is an inherently complex issue for both the deterministic approach and the stochastic approach. Since this study aims to compare these two types of computation, it is important to recognize the different methods used to combine stochastic reaction and diffusion [6-9, 44, 59, 69-71]:

1. First Passage-Time Kinetic Monte-Carlo method (FPKMC) [72-74] directly combines stochastic bi-molecular binding with Brownian diffusion using an exact, event-based approach. This algorithm takes into account excluded volume and crowding effects. Approximate implementations include the Green’s Function Reaction Dynamics method (GFRD) [75, 76]) and the Cellular Dynamics Simulator [77]. FPKMC has also been extended to the Doi volume reactivity model of bimolecular binding [60].
2. Discrete-time particle-based Brownian reaction dynamics (BRD) simulators (e.g. Smoldyn [78], MCell [79], GridCell [80]) can be viewed as approximations of the FPKMC/GFRD method. Since bi-molecular binding probability is not modeled exactly by a finite time-step method, convergence of results with time step size has to be carefully checked.
3. Course-grained stochastic simulation algorithm referred to as the Reaction-Diffusion Master Equation (RDME) method (e.g. SmartCell [81], MesoRD [82], URDME [83], STEPS [84], Lattice Microbes [85]) implement reactions *exactly* in elementary sub-volumes using the Gillespie stochastic simulation algorithm [86], assuming that reactants are well-mixed in each sub-volume. Diffusion is treated as an exchange reaction between neighboring voxels. Spatial resolution is limited by sub-volume size, which cannot be arbitrarily reduced without losing all reactions [59, 60, 87-89]. A convergent modification of this approach resolves the latter problem by allowing reactions between neighboring cells [90].

A large variety of hybrid methods have also been developed, for instance methods combining the advantages of RDME and BRD algorithms [27, 91, 92], as well as hybrid methods combining in various ways the stochastic and deterministic reaction-diffusion components [14-16, 30, 93-98]. Finally, for larger number of reactants, the Langevin approximation and formulations based on stochastic partial differential equations can be used [8, 71, 99-101].

For our comparative study, we will use Smoldyn [78], a particle-based BRD method, because of its flexibility, ease of use, and computational efficiency. This algorithm uses the Smoluchowski surface-reactivity model of bi-molecular reaction, with binding radii calibrated by approximate matching of the corresponding macroscopic mass-action reaction rates, making this method particularly well-suited for comparing with the deterministic mass-action simulation approach. This allows us to use the same reaction and diffusion parameters values in the stochastic and in the deterministic model of Ca^2+^ buffering and diffusion, without modification. It should be remembered however that the relationship between the macroscopic binding rates (“propensities”) and the underlying microscopic binding radius is a separate, nontrivial problem [8, 59, 60, 78, 88, 102-104]. Moreover, the functional form of mass-action bimolecular reaction rate is known to be violated at large densities [105, 106]. However, it is precisely the sum of all sources of quantitative discrepancy between straightforward mass-action and stochastic approaches that we want to investigate in this study.

To check the convergence of results with respect to the time step size, we repeated Smoldyn simulations for a range of time steps. In the results shown in Figs. 1-2, the time step size is set to Δ*t*=1 μs, while Δ*t*=0.2 μs was used for the case of larger Ca^2+^-sensor binding rates in Fig. 3. Simulation in Figs. 1-3 took over a month of total computation time with 12 simultaneous Smoldyn threads, while the corresponding deterministic simulations take less than one CPU-minute.

## III. RESULTS

### III.1 Spatially resolved 3D model

From a practical point of view, local fluctuations in Ca^2+^ concentration ultimately reveal themselves through the fluctuations in the latency to vesicle membrane fusion or other macroscopic Ca^2+^-dependent effects, which is the most important observable in the modeling of Ca^2+^-dependent exocytosis. Therefore, in our comparison of deterministic and stochastic modeling of vesicle release, we will focus on the latency between the opening of a Ca^2+^ channel, and the time of fusion of nearby vesicle(s). This latency is given by the first passage time (also known as the *waiting time* or *hitting time*) to the fully bound final “release” state *R* of the vesicle’s putative Ca^2+^ sensor [13, 14, 46-51]. We assume that the binding kinetics are described by Eqs. 3-5, inferred from the studies of the calyx of Held synapse [56, 57]. The qualitative conclusions of this work however do not depend on the type of model used for the sensor. In our simulations of 3D Ca^2+^ diffusion, buffering and binding, illustrated in Fig. 1, the sensor is located at a distance of 33nm away from the channel, within the channel’s “*nanodomain*”. The sensor position is marked by a square in the bottom-left corner of the diffusion volume in Fig. 1A,B: the two panels illustrate one time frame of the deterministic and stochastic simulation, respectively, as described in Methods and the figure caption.

In the full stochastic approach, each MCMC trial provides one sample of the Markovian sensor state transition time series shown in Fig. 1D, driven in turn by the stochastic Brownian motion of Ca^2+^ and buffer particles illustrated in Fig. 1B. The simulation is repeated in order to construct a histogram estimate of FPTD. In Fig. 1E, the resulting FPTD estimate is shown as a black curve, and was obtained using about 10^8^ Smoldyn iterations, each of which in turn comprises thousands of elementary MCMC iterations updating particle positions and binding states. In contrast, in the deterministic mean-field approach the FPTD is computed as the rate of change of the occupancy of the final absorbing state of the exocytosis sensor, obtained by numerically solving the coupled PDE-ODE reaction-diffusion problem (Eqs. 2-5). In Fig. 1E we directly compare FPTDs obtained using these two approaches, for the case when only *N*_Ca_=100 ions enter the model volume during a 1ms-long Ca^2+^ channel current pulse. In this first comparison we set the binding sink term *σ*_S_ in Eq. 2 to zero, ignoring the depletion of Ca^2+^ ions due to their binding to the sensor. Figure 1E reveals a surprisingly good agreement between stochastic and deterministic simulations of FPTD, despite this simplification. Note that no curve scaling is involved, or would be appropriate, in comparing the FPTD obtained using the two methods.

Part of the discrepancy between the two methods seen in Fig. 1E is explained by the depletion of five Ca^2+^ ions as they bind to the Ca^2+^ sensor, which is absent from a straightforward implementation of the mass-action approach that doesn’t include the feedback from the Ca^2+^-sensor binding onto the local Ca^2+^ concentration [24, 29]. To reveal the effect of this Ca^2+^ ion depletion, in Fig. 2 we repeat the numerical solution of Eqs. 2-5 with and without the Ca^2+^ sink term, given by Eqs. 6-7. Simulations are then repeated for several values of Ca^2+^ current strength corresponding to different number of total Ca^2+^ ions entering the volume during a 1ms-long channel opening, from *N*_Ca_=200 (Fig. 2A) down to *N*_Ca_=20 (Fig. 2D). Note that the depletion correction almost fully abolishes the discrepancy between stochastic and deterministic simulations for the case of 100 Ca^2+^ ions, but not for smaller number of ions. These results suggest that deterministic results become unreliable when only ∼40 or less Ca^2+^ ions enter the simulation volume (Figs. 2B-D).

While full parameter sensitivity analysis is prohibitive in view of the computational cost of the full stochastic approach, in Figure 3 we repeat our comparison with the binding rate increased by a factor of 10, corresponding to an unrealistically large sensor binding radius. To compensate for such a large Ca^2+^ binding rate, we also increased the distance from the Ca^2+^ channel to the release sensor, from 33nm to 83nm. As Fig. 3 shows, in this case the depletion correction (Eqs. 6-7) is critical for achieving good agreement between the two approaches even for a relatively large number of Ca^2+^ ions. Note also that the peak release occurs before the end of the current pulse, due to faster saturation of the Ca^2+^ sensor caused by the larger binding rate. However, despite significant difference in parameters used in Fig. 3 vs. Fig. 2, in both cases the discrepancy between stochastic and depletion-corrected mass-action simulations is apparent only when the total number of entering Ca^2+^ ions falls below about 40-50. The close agreement between the two approaches for such small total numbers of Ca^2+^ ions is *a priori* surprising, and is the focus of further analysis presented below.

We note that in instances when the stochastic FPTD appears smaller in amplitude compared to the deterministically computed FPTD, as is the case in Fig. 2B-D and Fig. 3D, the tail of the stochastically computed FPTD is much longer, since the cumulative binding probability always equals one, due to the closed domain with reflective boundary conditions.

To understand the close agreement between the results of 3D simulations obtained using the two approaches, we will now turn to a simplified model of this reaction-diffusion process.

### III.2 Analysis of a simplified two-compartment model

To gain an intuitive understanding of the factors affecting the relative accuracy of the mass-action approach, we will analyze a highly simplified model of Ca^2+^ diffusion and binding shown in Fig. 4, similar to the one analyzed by S.H. Weinberg [48] (see also [26, 52]). This reduced model consists of two well-mixed compartments of Ca^2+^ ions, with the Ca^2+^ sensor for exocytosis contained within the inner compartment. We assume that the sensor can bind two Ca^2+^ ions before triggering exocytosis, according to the reaction

**Figure 4.**
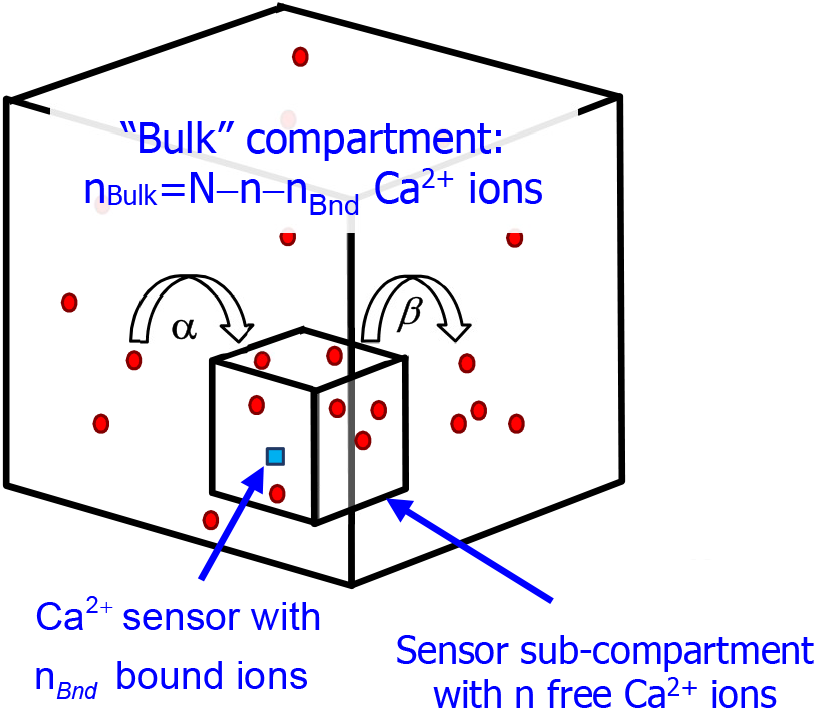
Two-compartment model of Ca^2+^ diffusion and sensor binding. The inner compartment contains *n* ions of Ca^2+^, which can bind to the Ca^2+^ sensor. The inner compartment exchanges ions with the bulk compartment with diffusive rates *α* and *β*. The total number of ions equals *N* and is conserved, therefore there are *n*_Bulk_=*N−n −n*_Bnd_ ions in the bulk compartments, where *n*_Bnd_ is the number of ions bound to the sensor, as described by Eq. 11. At initial time, all *N* particles are added to the bulk compartment.

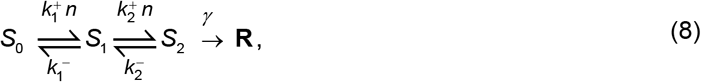

where *n* is the number of free Ca^2+^ ions in the sensor sub-compartment. For the sake of simplicity, the stoichiometric factors of 2 are absorbed into the definitions of the forward and backward binding rates, 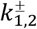 We set 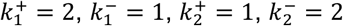.

Diffusion is represented as transitions between the two compartments, with forward and backward transitions rates equal to *α* and *β*, respectively:

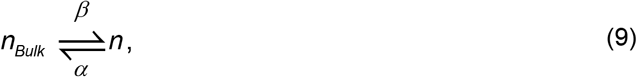

where the *n*_Bulk_ is the number of free Ca^2+^ ions in the bulk compartment, given by

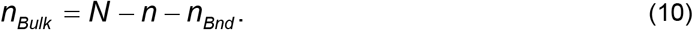

Here *N*=const is the total number of Ca^2+^ ions in both compartments, and *n*_Bnd_ is the number of sensor-bound Ca^2+^ ions, determined by the occupancies of all Ca^2+^-bound sensor states, indicated by angled brackets (for precise notation description, see Appendix, Eq. 16):

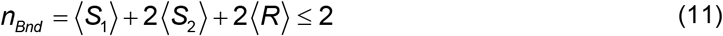

The ratio of diffusion rates *α* and *β* in Eq. 9 implicitly defines the ratio of the volumes of the two compartments. The total number of Ca^2+^ ions, *N*, is a constant parameter: initial condition is set by adding *N* ions to the bulk compartment. Therefore, initially there are no ions in the sensor compartment, nor bound to the sensor: *n*(0)=*n*_Bnd_(0)=0 (alternative initial conditions were also explored, but did not provide significant new insights). Even though Ca^2+^ buffering is not explicitly implemented in this simplified model, buffering can be viewed as being part of the exchange reaction with the bulk compartment described by Eq. 9, whereby buffer-bound Ca^2+^ ions are to be understood as belonging to the bulk (non-sensor) compartment.

The stochastic implementation of this model is given by the following continuous-time Markov chain, with (*n, S*_*k*_) and (*n, R*) denoting a state with *n* Ca^2+^ ions in the sensor sub-compartment, and the sensor in state *S*_*k*_ or in the final fusion state *R*:

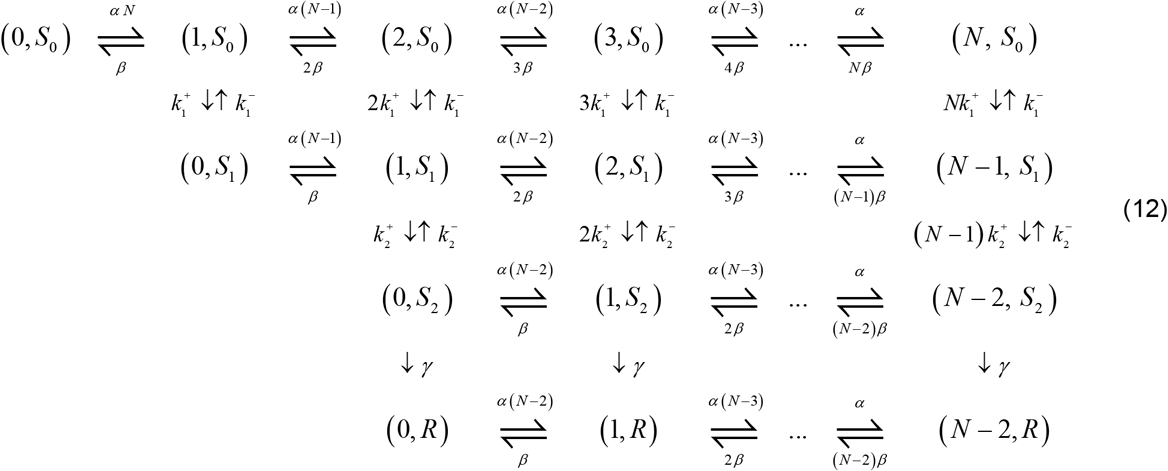

This model is almost equivalent to the one analyzed by S.H. Weinberg [48] (see Eq. 2.8 therein), except that we compare this model with a similar two-compartment deterministic representation of the same process, rather than its single-compartment reduction, allowing us to clearly separate the effects of binding-induced Ca^2+^ ion depletion from the impact of stochastic fluctuations. Another difference of our approach is that we consider the bulk compartment of a finite volume. Therefore, no particle number truncation is required, since the total number of Ca^2+^ ions is conserved and set equal to *N*.

The Markov Chain described by Eqs. 12 is readily converted to linear Chemical Master Equations (CME) describing the evolution of state probabilities, d**p**/dt = *W***p**, where vector **p** has 4*N*−1 states shown in Eq. 12. We note parenthetically that the number of states can be further reduced using sensor state conservation law and by collapsing together all (*n, R*) states. The CME system is shown explicitly in Eq. 15 of the Appendix. The CME system can in turn be converted (using appropriate summations) to the ODEs for the moments of the state variables, i.e. the occupancy probabilities of sensor states ⟨*S*_*k*_⟩ and ⟨*R*⟩, and the moments of the Ca^2+^ ion number *n* conditional on the state of the sensor, denoted as ⟨*n*^*m*^ |*S*_*k*_⟩ (see Appendix). The first five of these moment equations read:

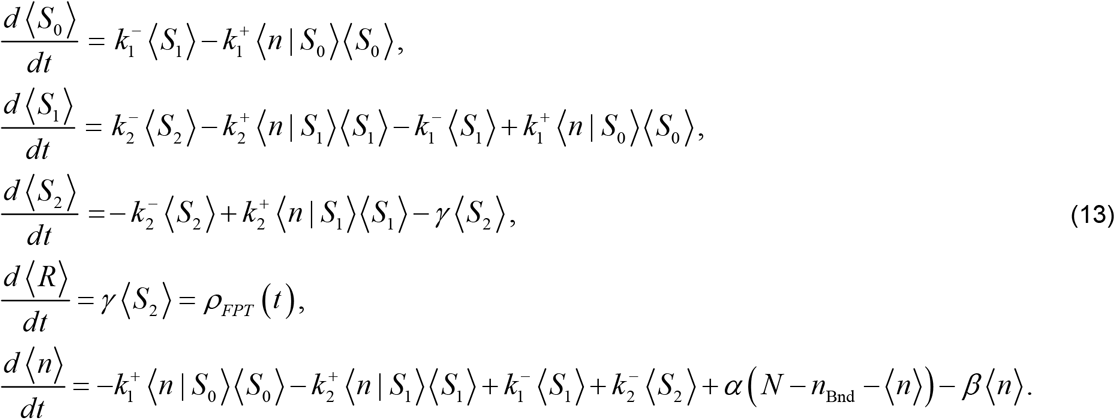

Here *n*_Bnd_ is given by Eq. 11 and denotes the total number of Ca^2+^ ions bound to the sensor, while *ρ*_FPT_ denotes the FPTD. The evolution equations for the first moments ⟨*n* | *S*_*k*_⟩ are not shown, but are easily derived from the CME system; they depend in a somewhat complex way on higher moments ⟨*n*^2^ | *S*_*k*_⟩. Since we consider a finite number of Ca^2+^ ions, the system of moments is closed and always solvable in closed form due to the linearity of the CME system.

Note that if the number of Ca^2+^ ions in the sensor compartment is independent of the sensor state, ⟨*n*|*S*_*k*_⟩=⟨*n*⟩, then Eq. 13 reduces to the deterministic, mean-field description of the same process,

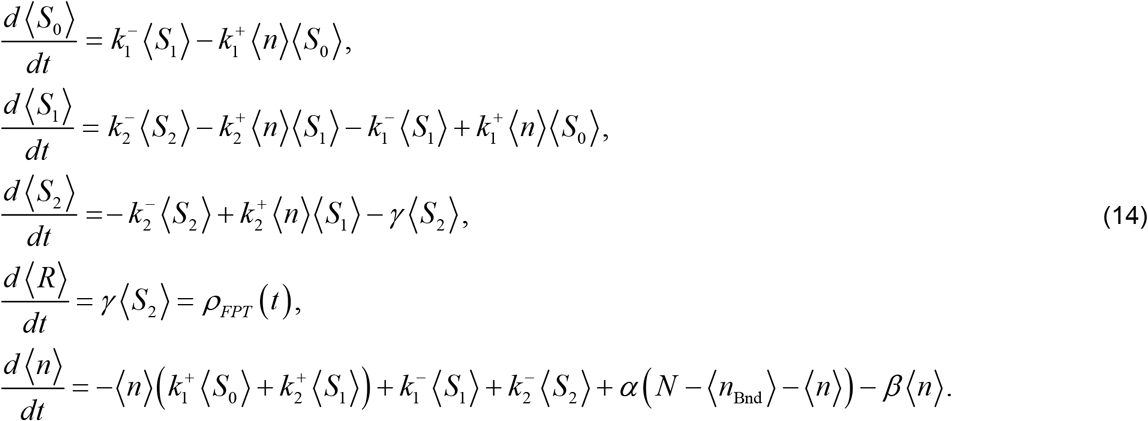

Thus, this mean-field description can be viewed as the simplest moment-closure of the stochastic description given by Eqs. 13 under the assumption of zero correlation between the state of the sensor and the number of ions in the sensor compartment. Therefore, the high variance in the local number of Ca^2+^ ions has little effect on FPTD computed using the mean-field approach, as long as the number of ions in the sensor sub-compartment is only weakly correlated with the binding state of the release sensor.

This intuition is confirmed by the results shown in Fig. 5, where we compare the exact solution of the stochastic two-compartment model described by Eqs. 12-13 with the solution of the mean-field description given by Eq. 14, for a small number of Ca^2+^ ions, namely *N*=4. The diffusion rates *α* and *β* are set to equal each other, to ensure that exactly one Ca^2+^ ion remains free (on average) inside the sensor sub-compartment upon equilibration, after 2 out of the 4 available Ca^2+^ ions are bound by the sensor. Figure 5 shows this comparison between stochastic and mean-field results for three distinct ratios between diffusion and reaction rates: *τ*_diff_ */τ*_R_=100 (Fig. 5A1-C1), *τ*_diff_ */τ*_R_ = 3 (Fig. 5A2-C2) and *τ*_diff_*/τ*_R_ =0.01 (Fig. 5A3-C3). This ratio is controlled by varying the diffusion rate *α*=*β*=1/*τ*_diff_, while keeping the reaction rates constant (*τ*_R_=1/*k*_1_^−^=1/*k*_2_^+^=1). As Figure 5 demonstrates, the difference between the FPTD distributions obtained using the mean-field and stochastic approaches can be surprisingly small even for *N*=4∼O(1). In agreement with the arguments above, the discrepancy between the FPTD obtained using the two approaches is the smallest for the case when the absolute correlation between the sensor state and the number of Ca^2+^ ions in the sensor sub-compartment (Fig. 5C1,C2,C3) is also the smallest, which happens for *τ*_diff_*/τ*_R_ ≈ 3. In this case the diffusive fluctuations are sufficiently fast relative to the reaction, partially “washing out” the correlations between the sensor state and the number of Ca^2+^ ions in the sensor compartment. The correlations are quantified in Fig. 5C1,C2,C3 in terms of the ratio of conditional and unconditional moments of the Ca^2+^ ion number inside the sensor compartment; the latter are explicitly shown in Fig. 5B1,B2,B3.

**Figure 5.**
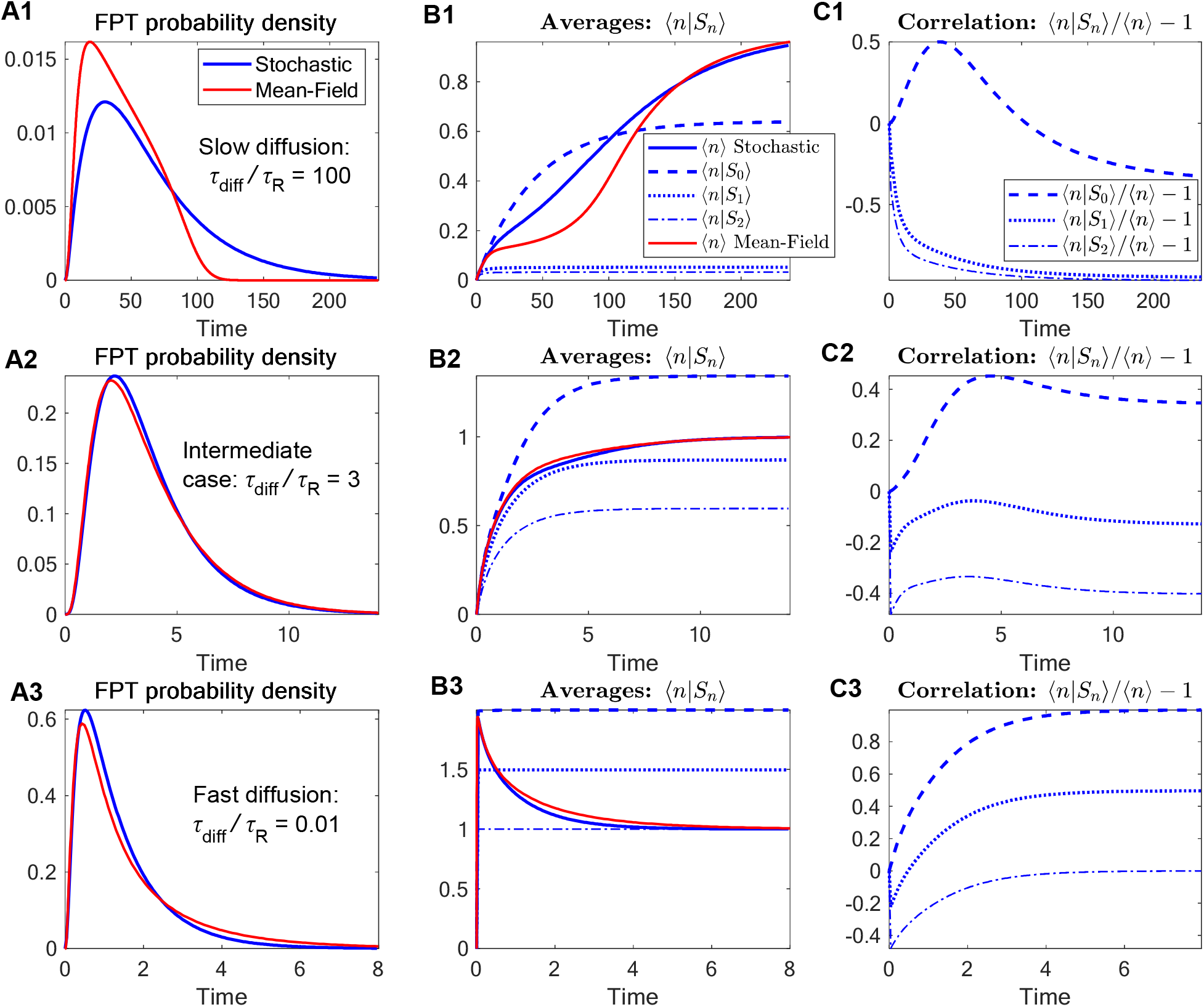
Comparison between mass-action and stochastic simulation of the two-compartment model in Fig. 4, for different ratios between diffusion and reaction time scales: *τ*_diff_ */τ*_R_=100 (**A1-C1**), *τ*_diff_ */τ*_R_=3 (**A2-C2**), and *τ*_diff_*/τ*_R_=0.01 (**A3-C3**). This ratio is controlled by varying the diffusion rate *α=β*=1*/τ*_diff_, while keeping reaction rates constant. (**A1**,**A2**,**A3**) First-passage time to full binding probability density (FPTD). (**B1**,**B2**,**B3**) Average number of Ca^2+^ ions in the sensor compartment. For the stochastic simulation, also shown are the average number of Ca^2+^ ions conditional on the sensor state, ⟨*n*_Ca_|*S*_*k*_⟩. Note that the number of ions in the sensor compartment approaches *n*=1 in all cases, since the total number of Ca^2+^ ions is *N*=4. (**C1**,**C2**,**C3**) Correlation between the number of Ca^2+^ ions in the sensor compartment and the sensor state, given by the ratio of conditional and unconditional expectations ⟨*n*_Ca_|S_*k*_⟩/⟨*n*_Ca_⟩ (*k*=0,1,2).

The dependence of correlations on the *τ*_diff_ */τ*_R_ ratio can be quite non-trivial, but intuitive. For example, in the case of slow diffusion (Fig. 5C1), the largest correlations (in absolute value) are negative, ⟨*n*|*S*_1,2_⟩/⟨*n*⟩−1→ −1, since ⟨*n* |*S*_1,2_ ⟩<<1. This is because there is only a small probability of any ions remaining in the sensor sub-compartment when at least one ion is bound to the sensor: the fact that the sensor is still not fully bound indicates that the remaining ions are most likely outside of the sensor compartment. In contrast, for fast diffusion (Fig. 5C3), the largest correlation approaches ⟨*n* | *S*_0_⟩ / ⟨*n* ⟩ − 1 → 1, or ⟨*n* | *S*_0_ ⟩ ≈ 2⟨*n* ⟩: when the sensor is unbound, there are on average *n*=*N*/2=2 Ca^2+^ ions inside the sensor compartment, which is twice as many as there will remain upon full sensor binding. Recall that we set *α* =*β*, so ions quickly equipartition between the sensor compartment and the bulk in the limit of fast diffusion.

Interestingly, results shown in Fig. 5C1,C2,C3 suggest that the dependence of the maximal correlation size on the diffusive time scale is non-monotonic. This is confirmed by a more detailed comparison shown in Figure 6 as a function of the diffusive time scale, *τ*_diff_=1/*β* (assuming again a constant reaction rate and *α* =*β*, for the sake of simplicity).

**Figure 6.**
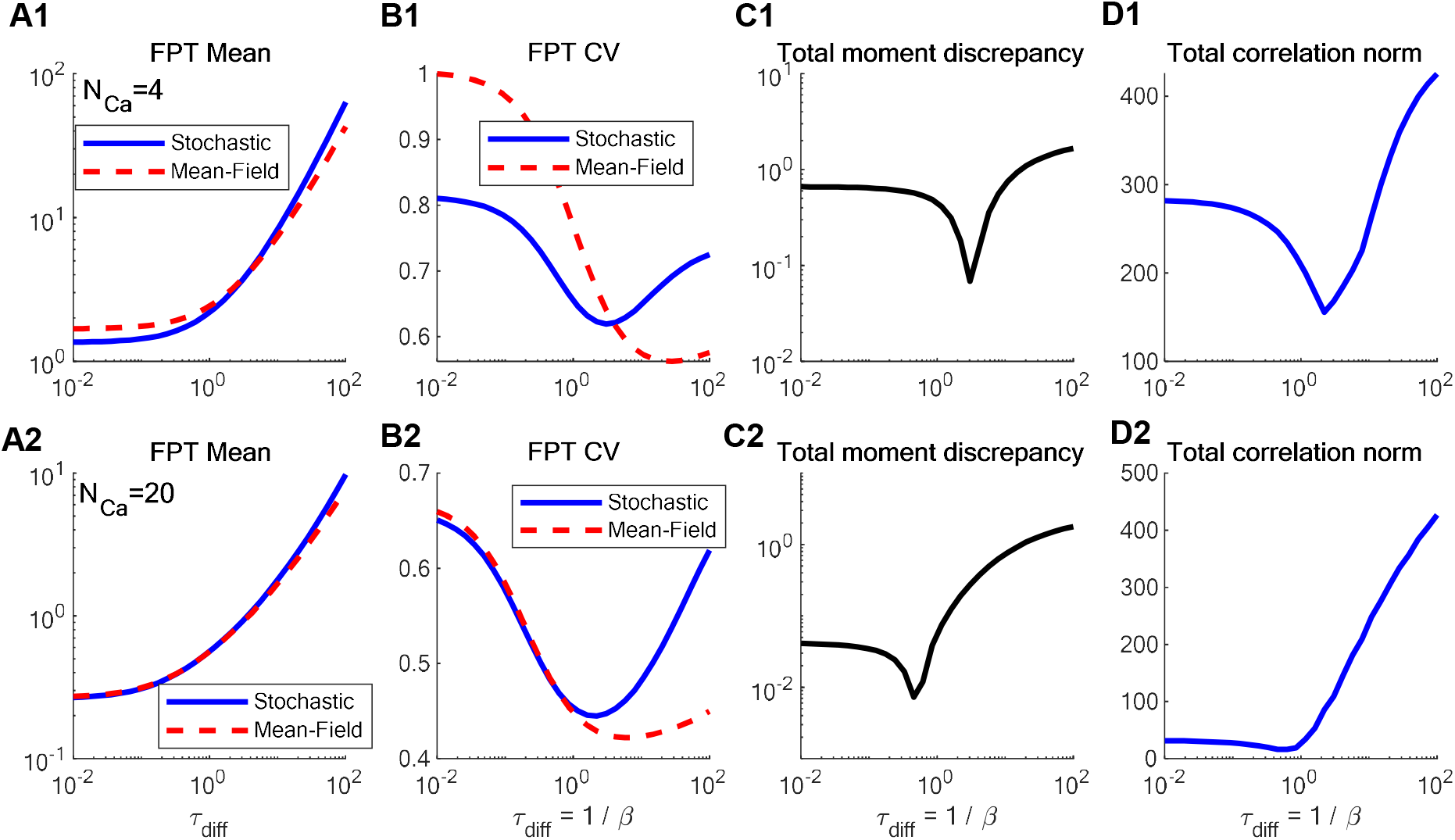
Comparison of the correlations and the central moments of FPTD obtained using deterministic mean-field vs stochastic approaches, as a function of diffusive time constant τ_diff_=1/β, for two values of total particle number, *N*=4 (A1-D1) and *N*=20 (A2-D2). (**A**1, **A2**): FPT average, (**B1, B2**): coefficient of variance of FPT, (**C1, C2**): total normalized discrepancy of first 3 central moments of FPTD between stochastic and deterministic simulations. (**D1, D2**) sum of absolute correlations ⟨*n* | *S*_0_⟩, ⟨*n* | *S*_1_⟩ and ⟨*n* | *S*_2_⟩ in the stochastic model. Diffusive rates satisfy α/β=1 for all simulation conditions, ensuring that the average number of ions in the sensor compartment upon full sensor binding is *n*=1 in (A1-D1) and *n*=9 in (A2-D2).

Intuition suggests that in the limit *τ*_diff_→∞, diffusion is very slow on the time scale of the reaction, and therefore the correlation between the sensor state and the number of Ca^2+^ ions in the sensor compartment imposed by the Ca^2+^-binding reaction is strong, resulting in a large error of the mean-field reduction of the problem. Figure 6 confirms this intuition, showing that this discrepancy between stochastic and deterministic computation of FPTD grows as the diffusion rate decreases. However, Fig. 6D1,D2 also reveals that the dependence of FPTD discrepancy on diffusive time scale is non-trivial, reaching a minimum for certain fixed relationship between the rates of diffusion and reaction. Therefore, correlations between the sensor state and the number of Ca^2+^ ions in the sensor sub-compartment do not disappear as *τ*_diff_→0. In this limit, the numbers of Ca^2+^ ions in the bulk compartment and the sensor sub-compartment are in instantaneous equilibrium with each other, so a Ca^2+^ binding reaction leads to an immediate reduction of the latter. Thus, in the case of very fast diffusion, Ca^2+^-binding reactions create strong negative correlations between the sensor state and the number of Ca^2+^ ions, which is lower in absolute magnitude than in the case of very slow diffusion, but still significant. Therefore, in the compartmental model, the correlations and the discrepancy between mean-field and stochastic computations of FPTD reach its minimum values for certain intermediate level of the ratio between the diffusion and reaction time scales. However, for larger number of Ca^2+^ ions, this non-monotonic relationship is less significant, as shown in Fig. 6A2-D2: the discrepancy between the two approaches is in general more pronounced in the limit *τ*_diff_→∞.

Another way to understand the growing inaccuracy of the mean-field approach as *τ*_diff_→0 is that the two-compartment mean-field description given by Eq. 14 does not approach the physically correct model in this limit, since it does not take into account the immediate reduction of the free Ca^2+^ ion number *n* upon binding. In other words, in the limit *τ*_diff_→0 (*α* → ∞, *β* → ∞) the number of ions in the sensor compartment is no longer an independent variable, and is directly determined by the sensor’s binding state: *n* → *α*(*N* − *n*_Bnd_)/(*α* + *β*). Unlike the mean-field description, the stochastic description given by Eq. 13 remains meaningful for any values of rate parameters.

## IV. Discussion

Our main finding is that the discrepancy between stochastic and deterministic simulation of Ca^2+^ diffusion and bi-molecular binding/unbinding reactions can in certain cases be much smaller than expected from naïve intuition (i.e., naïve application of 1/√*N*_ca_ scaling). This result has very practical significance, since stochastic Ca^2+^ channel gating (and more generally, fluctuating Ca^2+^ current) is computationally inexpensive to simulate and combine with deterministic mass-action reaction-diffusion equations, providing an efficient hybrid method for the modeling of Ca^2+^-dependent phenomena [7, 14, 30].

However, we want to emphasize once again that the accuracy of such an approach was shown to be greatly reduced in the presence of positive feedback on Ca^2+^ influx provided by CICR [19-28, 107]. We should point out that recent studies suggest that CICR does contribute to both pre- and post-synaptic processes [108-112]. Still, it is useful and important to explore the impact of fluctuations on each part of the biochemical pathways to vesicle release, including those not involving CICR.

To complement prior work on the role of stochastic fluctuations in Ca^2+^-dependent mechanisms, we considered a maximally reduced model to focus on two sources of stochasticity downstream of Ca^2+^ influx, namely the diffusive fluctuations in Ca^2+^ concentration, and the fluctuations due to Ca^2+^ binding and unbinding reactions. Further, following several prior studies [13, 14, 46-51], we focused on the modeling of FPTD, which can be considered as the true “output observable” whose uncertainty combines all upstream sources of stochasticity in this biochemical process. Considering such maximally simplified, reduced model allowed us to study more directly the interplay between fluctuations due to diffusion and reaction in determining the final fluctuations in the Ca^2+^ sensor binding time.

Simulations of the reduced spatially-resolved model revealed that the distribution of FPT to full binding of the exocytotic Ca^2+^ sensor is accurately predicted by the mass-action / mean-field approach, as long as the number of Ca^2+^ ions is above about 50 (Figs. 1-3). This rough estimate of the threshold of accuracy seems to be relatively stable with respect to several parameters of the model, as seen for instance by comparing Figs. 2 and 3 that differ considerably in several critical model parameters. We note that this is a surprising result, given the presence of multiple bimolecular reactions, since the associated nonlinearities are expected to amplify the impact of fluctuations in the local Ca^2+^ ion concentration [44]. Intuition may suggest that buffering “washes out” stochastic fluctuations in the local Ca^2+^ ion numbers, reducing the contribution of such fluctuations to the fluctuation in FPT. Interestingly, it has been shown [52] that exactly the opposite is the case: namely, mobile buffers typically *increase* stochastic fluctuation amplitude. Therefore, another explanation for the close agreement between the two approaches is needed.

Our analysis of the simple two-compartment model suggests such an explanation. Namely, the inaccuracy of the deterministic description of FPTD is primarily determined by the correlations between fluctuations in the reactant molecule numbers, rather than the size of these fluctuations. This in turn follows from the fact that all reactions relevant to the Ca^2+^ buffering and sensing are *hetero-species* rather than *homo-species*: reactions occur between molecules of different types [59]. In this case the mean-field description of the Ca^2+^ sensor approximates its stochastic description, under the simplifying assumption of zero correlations (cf. Eqs. 13-14). This correspondence would break down had Ca^2+^ sensing involved simultaneous binding of two Ca^2+^ ions to the exocytosis sensor, as opposed to a sequence of two binding reactions. This scenario would correspond to a larger ratio of between reaction and diffusion rates, which is expected to amplify the discrepancy between deterministic and stochastic approaches. These results are quite general and apply to the modeling of a wide class of biochemical cell processes. The significance of the relative magnitudes of reaction and diffusion rates have also been pointed out by prior studies (see e.g. [16, 23, 24, 38, 48]).

It is important to emphasize that the partial decorrelation between local Ca^2+^ fluctuations and the sensor state fluctuations does not imply a small uncertainty in FPT: in fact, FPTD is still quite “wide” (i.e. has a large coefficient of variance) even in cases where the mean-field results achieve significant accuracy (see Figs. 1-3). We note that this finding is similar to, but somewhat distinct from the concept of stochastic shielding, whereby stochastic fluctuations of upstream reactions in a given biochemical pathway are effectively shielded from fluctuations of observable variables (such as open states of an ion channel), which are more sensitive to fluctuations in downstream parts of the pathway [113, 114].

The above-mentioned threshold of *N*_Ca_ =50 inferred from our spatial simulations shown in Figs, 1-3 is not wholly satisfying, considering that our analysis of the reduced two-compartment model illustrates that good agreement between stochastic and deterministic (mean-field) estimates of FPTD can be achieved even for *N*_Ca_∼O(1). Further, relating the discrepancy between deterministic and stochastic simulations to the ratio of diffusion and reaction rates is conceptually problematic, since bi-molecular binding reactions are limited by diffusion, precluding truly independent variation of reaction and diffusion rates. Therefore, it would be crucial to bridge the conceptual gap between compartmental and fully spatially resolved models, while retaining the ability for rigorous analysis. The most promising approach in this direction is to use recent simplified models of Ca^2+^ diffusion, buffering and binding that allow closed-form computation of FPTD [49-51]. Although the latter studies involve certain simplifications, particularly in their reduced descriptions of Ca^2+^ buffering, it would be promising to examine the discrepancy between stochastic and deterministic FPTD obtained in the framework of such simplified models.

It is possible that some part of the discrepancy between deterministic and stochastic methods that we observed with smaller number of Ca^2+^ ions could arise from the difference in the treatment of the exocytosis sensor’s Ca^2+^ binding used in the two approaches, as described in Methods. However, we believe it is important to compare the most straightforward and easily implementable approaches, which are most often used in practice. Further study will confirm whether the discrepancy between the two approaches is significantly affected by the differences in the Ca^2+^ sensor binding implementation. However, in general, single-molecule scale effects can only be precisely computed using stochastic methods like FPKMC/GFRD [60, 72-77], or even more detailed molecular dynamics simulations [103].

## Acknowledgements

The Author acknowledges useful discussions with A. Donev (NYU) and M.G. Pedersen (University of Padova). This work was supported in part by NSF grant DMS-1517085.

## APPENDIX

### CME system for the two-compartment model

Let 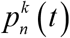 denote the probabilities of Markovian states in Eq. 12, corresponding to *n* (*n*=0..*N*) Ca^2+^ ions in the sensor sub-compartment while the sensor is in state *S*_k_, *k*=0..3, with *k=*3 indexing the terminal fused (release) state *R* in the sensor binding reaction, Eq. 8. Then the Markov Chain shown in Eq. 12 corresponds to the following Chemical Master Equation (CME) system (Kolmogorov forward equations):

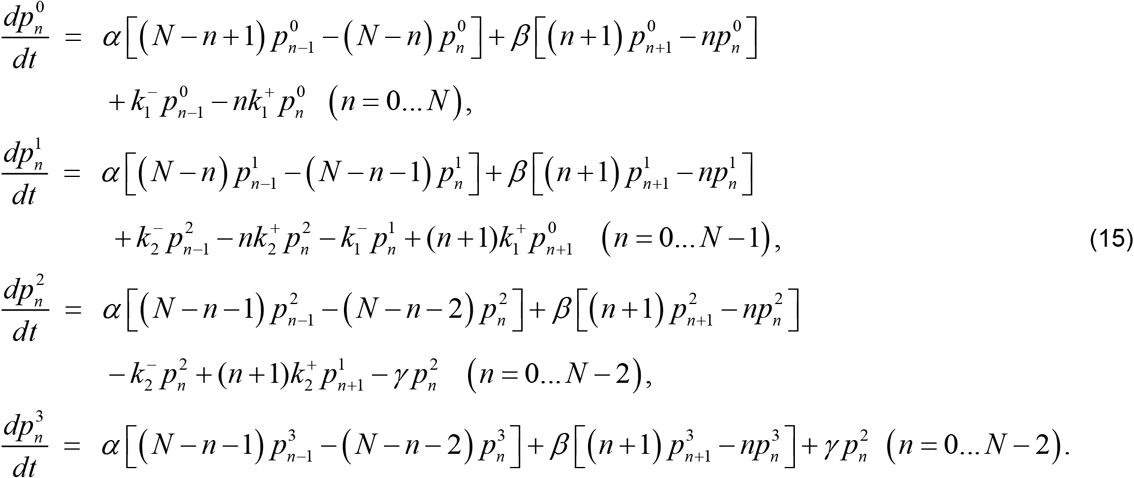

This is a linear system of form d**p**/dt = *W***p**, where the components of probability vector **p** correspond to the 4*N*−1 states shown in Eq. 12, and the elements of Markov transition matrix *W* are constant rates indicated in Eqs. 12, 15. Therefore, it is readily integrated in closed-form: **p**(*t*) = exp[*W t*] **p**(0). Since our quantity of interest is FPTD, given by the rate of transition to the final absorbing (release) state *R*, all states (*n, R*) with probabilities 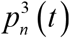 could be collapsed onto a single absorbing state by summing over *n*. Further, one more variable could be eliminated using the conservation law for the sum of all states of the sensor. However, we keep all states in Eqs. 12, 15, for the sake of clarity.

This CME system can be converted to an ODE system of the same dimensionality describing the moments of the state variables, i.e. the occupancy probabilities of sensor states ⟨*S*_*k*_⟩ and ⟨*R*⟩, and the moments of *n*, which depend on the state of the sensor, and denoted as ⟨*n*^*m*^, *S*_*k*_⟩:

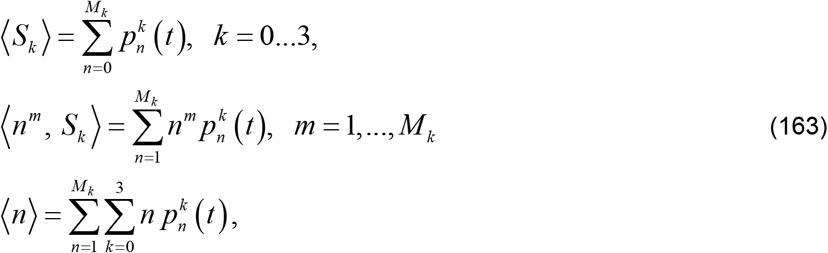

where *M*_*k*_ = *N* − min (*k*, 2) is the number of free (unbound) Ca^2+^ ions contained in both compartments when the sensor is in state *k*. Introducing conditional moments through the standard definition ⟨ *n*^*m*^, *S*_*k*_ ⟩ = ⟨*n*^*m*^ | *S*_*k*_ ⟩ ⟨*S*_*k*_ ⟩, and performing appropriate summations of Eqs. 15 and algebraic simplifications, one obtains the moment system, part of which is shown in Eq. 13. The initial condition corresponding to Figs. 5, 6 is given by 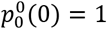, with all other probabilities satisfying 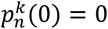.

